# Fecal Hormone Metabolites Concentrations Collected Longitudinally Confirm Spontaneous Polyestry in Mt. Graham Red Squirrels (*Tamiasciurus fremonti grahamensis*)

**DOI:** 10.64898/2026.05.29.728663

**Authors:** Stuart A. Wells, John L. Koprowski

## Abstract

The Mount Graham red squirrel (*Tamiasciurus fremonti grahamensis*, MGRS) is a federally endangered subspecies endemic to southeastern Arizona. Despite the establishment of *ex situ* conservation programs, reproduction outside the wild has not occurred. We monitored fecal hormone metabolites (FHM) to noninvasively assess estrous cycling and reproductive hormone dynamics in *ex situ*-managed MGRS. We compared *ex situ* findings with two *in situ* (wild) populations. We tracked longitudinal changes in estradiol and progesterone metabolites in three *ex situ* managed females and evaluated whether ovulation was spontaneous or induced. We detected 22 ovulation events over two years, with timing and regularity consistent with spontaneous polyestrous cycling. Comparative hormone analyses revealed appreciably lower progesterone and corticosteroid metabolite levels in *ex situ* individuals, whereas estradiol levels were comparable to those of wild counterparts. These findings suggest that maintaining environmental, and social conditions, e.g., lighting, temperature, and conspecific proximity, in *ex situ* settings is critical and could influence reproductive success. By confirming spontaneous ovulatory cycling in MGRS and demonstrating the utility of FHM as a noninvasive reproductive assessment tool, this study advances *ex situ* breeding efforts and provides a framework for reproductive monitoring in other endangered taxa.

## Introduction

This planet is currently experiencing its sixth major extinction event, driven by anthropogenic impacts[1–3]. The loss of terrestrial habitat and other anthropogenic factors has resulted in the extinction of nearly 200 vertebrate species since 1900, including at least 35 mammals [1–3]. As more species become threatened or endangered, it is increasingly important for wildlife conservation practitioners to collaborate closely with *ex situ* species management programs, such as zoos[4]. Such collaborations can guide *in situ* conservation, provide opportunities to support existing populations, and inform *ex situ* species managers about the critical components needed to give animals the best chance of surviving and thriving once released into the wild[4,5].

One of the most encouraging developments in *ex situ* managed wildlife conservation is support for species propagation, specifically for reintroduction into the wild [6]. The vital role of *ex situ* managed species propagation has advanced in recent years, especially as zoos have applied their unique expertise in developing *ex situ* species reproduction management with multiple taxa [7,8], [9], [10], [11], [12]. Advancements in animal husbandry and reproductive technologies provide essential tools for the overall sustainability of endangered species, both *ex situ* and *in situ*. Zoos focus on maintaining genetically viable collections, and *ex situ* zoo management expertise has led to the development and application of small-population genetic management tools to mitigate the loss of genetic viability in populations under managed care [13], [14], [15]. Zoological institutions also actively participate in wildlife species recovery through reintroductions and support habitat protection and restoration [4,16]. Zoo-based reintroduction programs (i.e., conservation breeding programs) provide effective and beneficial species conservation efforts rather than merely an alternative to *in situ* conservation action [11], [12]. Zoos have contributed considerably to the conservation of endangered species [17], [18], [12]. However, more work is needed to increase our understanding of the reproductive physiology and behavior of endangered species to mitigate the challenge of developing effective reproductive strategies, especially for animals held in *ex situ* settings and whose offspring are intended for release into the wild [4,17,19].

Zoos and other *ex situ* managed programs also promote innovative and emerging applications of assisted reproductive technologies for wildlife propagation. Assisted reproductive technologies (ARTs) encompass various biotechnologies, including artificial insemination and *in vitro* fertilization, as well as more complex techniques such as embryo transfer, somatic cell nuclear transfer (cloning), and sex determination procedures [20], [21], [22]. Many of these techniques, however, require invasive measuring of circulating reproductive hormones through blood draws and serum analysis. Restraining or anesthetizing animals for such procedures can affect hormone levels and lead to inconsistent results[23], [24]. The application of noninvasive collection methods, i.e., urine or fecal sampling [23], [25], can help to mitigate these adverse behavioral and physiological effects.

Fecal hormone metabolite analysis (FHM) is a powerful noninvasive tool for measuring metabolite concentrations resulting from circulating hormones, including estradiol, progesterone, testosterone, and corticosteroids [25]. Fecal samples can be collected after an animal is no longer present or in a way that does not disrupt its behavior (e.g., bedding, substrate) [26],[20],[27]. Measuring FHM is refined from assays developed for specific taxa [21] and can be applied broadly. Applying FHM techniques can also ascertain the physiological state of animals [28]. FHM enables the collection of longitudinal hormonal data from *ex situ* managed animals, resulting in continuous, and consequently, more robust, data analysis [29]. Monitoring through FHM has provided valuable insights into animal well-being by examining hormonal responses to social and environmental conditions [30], [31]. For example, FHM could be used to track ovulation events, which are physiologically indicated by a rise in estradiol metabolite concentration associated with increased progesterone levels [24], [25].

A noninvasive, longitudinal approach that reduces animal stress due to handling and does not affect circulating hormone levels could yield more accurate estimates of hormonal changes over time [23]. Due to their federal endangered status, their chronically low numbers in the wild, and the lack of successful *ex situ* reproduction, the Mount Graham red squirrel (*Tamiasciurus fremonti grahamensis*, MGRS) provides an ideal model for applying FHM technologies to inform and guide the development of *ex situ* conservation breeding programs.

It is important to use noninvasive reproductive assessments, such as FHM, because invasive or even minimally invasive stressors can affect circulating hormones and alter sociality and reproductive behavior over time, which are essential for reproductive success upon reintroduction into the wild [17], [23,32–34]. In general, mammals exhibit two modes of ovulation: induced ovulation, where ovulation (release of ovum) is dependent on copulation or other external stimuli (as in lions, ferrets, rabbits), and spontaneous ovulation, which occurs in the absence of copulation or specific external stimuli, most common in wild canids and domestic animals. The work conducted here represents only the second use of FHM applied to characterize reproductive hormone changes in a North American red squirrel, with Santymire et al. [35] being the first.

We were interested in determining whether MGRS exhibit spontaneous rather than induced ovulation cycles, as indicated by an increase in estradiol metabolites above 1 standard deviation, immediately followed by an increase in progesterone (i.e., on the same day) that lasts at least one day, occurring in the absence of copulation[36]. We were also interested in assessing whether FHM levels would differ appreciably between *ex situ* managed and *in situ* red squirrels. These results could help refine an *ex situ* breeding management strategy, increase the likelihood of successful reproduction of this endangered species, and provide a conservation framework for other species.

## Methods

### Study system

The MGRS [37] is the southernmost subspecies of the North American red squirrel [38], [39],[37]. Endemic to mature to old-growth mixed conifer ecotones and spruce-fir forests in the upper elevations of insular Mount Graham (i.e., Pinaleño Mountains) in southeastern Arizona [40], MGRS has remained reproductively isolated since the Pleistocene, approximately 10,000 years ago [41]. Due to a drastic population decline, primarily caused by habitat loss but also by wildfire and insect infestation, which impacted forage and refugia [42], [43], the MGRS was listed as Endangered under the US Endangered Species Act in 1987 [40]. The Mt. Graham Red Squirrel Recovery Plan [44] recommended developing an *ex situ* breeding program as a conservation strategy, given the continued decline of the wild population and other threats to extinction.

The MGRS has been studied extensively in the wild with much known about their sociality, habitat preferences, and resource requirements [45], [43,46,47], [48,49], [50], [51]. The MGRS is polygamous, and both sexes are intensely territorial and defend their territories, larder hoards. (termed “middens,”) from intruders [47], [45,52] [49], [46,53]. This intense territoriality relaxes when males detect and seek out receptive females to mate with during the breeding season, which spans approximately from February to September [37,54], [49], [45], [46]. However, the period of receptivity was believed to occur for only one day a year, within a four to six-hour window [55], [45], [40]. Although the MGRS have been in zoo-managed care [5], [35], they have yet to reproduce outside the wild, highlighting the importance of developing a consistent propagation and release program that produces animals suitable for release into the wild. A recent study of *ex situ* managed MGRS [35] revealed that MGRS exhibit polyestry, suggesting that breeding receptiveness may be temporally broader than previously suspected. The study conducted by Santymire et al. [35] employed FHM techniques to measure reproductive hormones, vulvar swelling, and vaginal lavage samples to detect cornified epithelial cells. Obtaining lavage samples in this manner necessitates some degree of invasiveness and animal acclimation to the procedure, which could impact circulating hormone levels due to the handler’s proximity to the squirrel [55], [20]. The vaginal lavage procedure, a process that has no comparable *in situ* occurrence, could also impact sociality and behavior over time [56], which could have unforseen consequences on reintroduction[17]. An entirely noninvasive design to assess hormone metabolite concentrations as conducted in this study, when effectively combined with reproductive behavioral indicators could mitigate stress-related hormone biases, and provide important insights to assist in developing an *ex situ* breeding strategy[57]. Additionally, once behavioral indicies for the onset of reproductve rediness are established, the need for ongoing FHM monitoring can be reduced or eliminated entirely.

Through collaboration with the US Fish and Wildlife Service (USFWS), the Arizona Center for Nature Conservation/Phoenix Zoo (ACNC) acquired 3 male and 3 female (3.3) squirrels between 2011 and 2014. We conducted this work under USFWS permit #TED1832-Z and in accordance with the University of Arizona’s Institutional Animal Care and Use Committee (IACUC) guidelines and animal husbandry guidelines developed specifically for MGRS. We housed the squirrels at ACNC’s Arthur L. and Elaine V. Johnson Native Species Conservation Center. We developed and applied animal care guidelines for this, with refinements made as more insights into best management practices became available[5]. Each squirrel was housed separately within a specially designed 1.5 x 1.5 x 1m enclosure due to their territorial behavior. The enclosures incorporated important MGRS territory demarcation features [58], including two attached nesting areas (76.2 x 30 x 30 cm) at opposite sides, which allowed squirrels to freely select which to use for nesting or as a larder (Fig 1). Each attached nest box was divided into two equal compartments by a removable internal gate.

**Fig 1.**
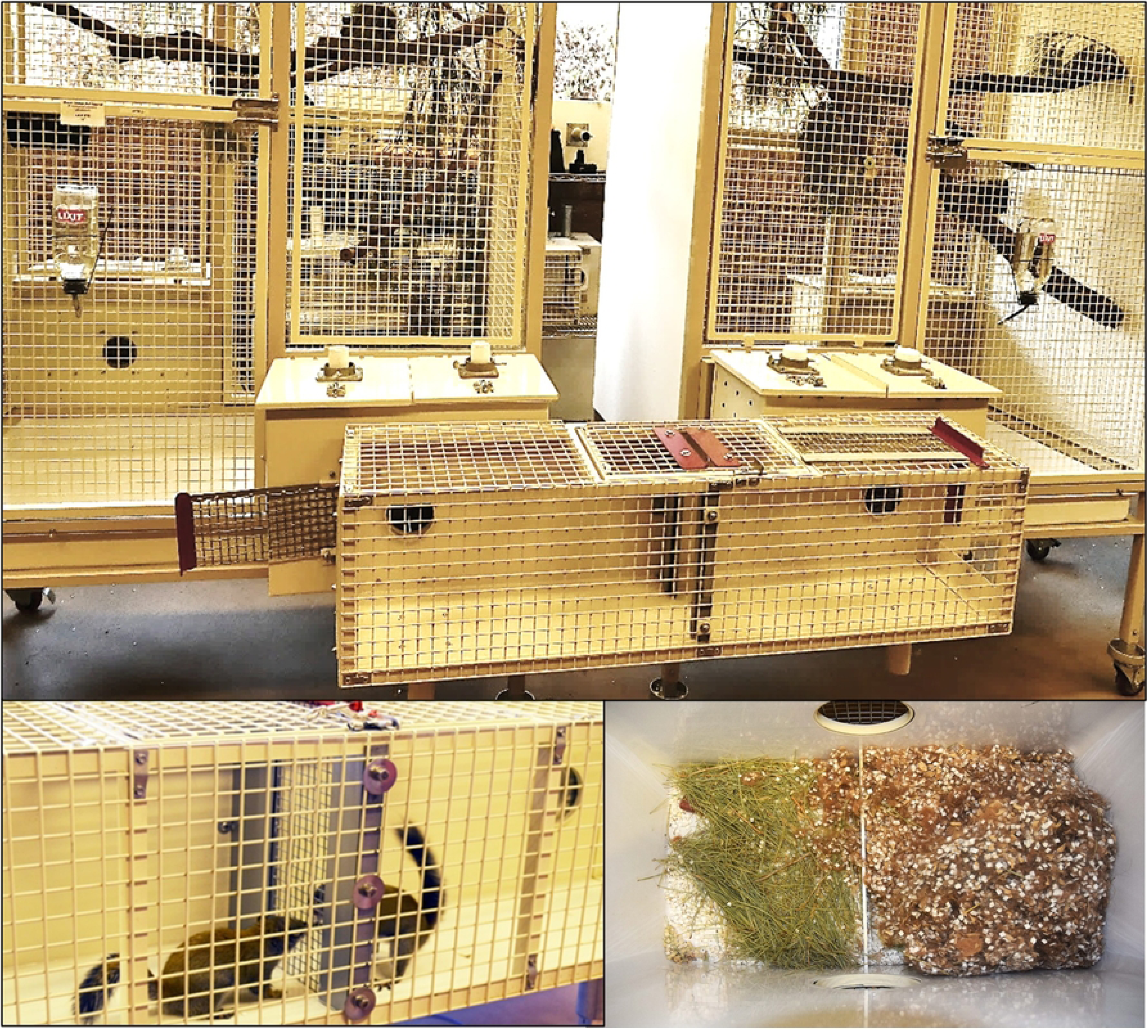
View of enclosure, nest box, and animals in attached cage, used for Mount Graham red squirrels (*Tamiasciurus fremonti grahmensis),* in Phoenix, Arizona. Nest boxes include AlphaDri™ substrate, pine needles. and shed mammal hair bedding (the squirrels incorporate the AlphaDri™ into the shed hair)

Each squirrel received 166g of diet per week, totaling 180 kcal. We provided the squirrels with an average of 24g of diet daily, divided into three feedings. Their diet was based on a nutritionally balanced formula developed in conjunction with a contracted professional nutritionist. Diet was dispersed throughout the enclosure to encourage natural foraging behavior. We equipped each enclosure with a 60ml glass water bottle with a bead-regulated sip nozzle.

Each day, we measured and recorded the squirrel’s water consumption, and we cleaned and sanitized the water containers. We maintained the squirrel’s environment at 18.3 ± 1 °C (65 ± 2 °F). The squirrels were exposed to natural light and to a seasonally adjusted lighting cycle consistent with the seasonal light patterns on Mount Graham(32.7017° N, 109.8715° W).

### Sample collection and FHM processing

From April 3, 2015, to February 28, 2017, fecal samples were collected daily. The samples were collected from the enclosure floor after the squirrel was closed within its nest box during routine daily care. To compare *ex situ* results with wild values, samples were also collected ≥3 per individual *in situ* from the Mount Graham population (MG) and the nearest red squirrel population (*T. f. fremonti*) in the White Mountains, Arizona (WM). Field sampling methods are described elsewhere and occurred throughout the year[59]. The storage, extraction, and processing of fecal corticosteroid metabolites, testosterone, progesterone, and estradiol were performed at the Lincoln Park Zoo’s Davee Center for Epidemiology. Specifically, this study employed Enzyme Immunoassay (EIA), a highly sensitive, accurate, and applicable technique for measuring FHM, thereby obviating the need for radioactive substances or parallel techniques [60].

### Defining and identifying ovulation events

We performed all analyses in R version 4.1.1 [49]. A mammalian spontaneous ovulation event occurs following a physiological increase in estradiol and progesterone [61,62], [63]. Our central question is whether the waiting times between ovulation events follow a periodic spontaneous (spontaneously initiated) distribution or an induced (externally stimulated) distribution [53]. To explore ovulation events in MGRS, we first identified local peaks in the hormone concentration curves of *ex situ* females. Then, we identified candidate “ovulation events” based on whether hormone peaks were aligned chronologically. Algorithmically, we defined an event as follows: where Δ days is the time difference between peaks (in days), and x is a positive integer representing the number of days between peaks. The default was to set x = 0, “same-day” peaks. We parameterized a function with a lag argument in an R script to allow a 1-2-day flexible window if necessary. We set cutoffs for estradiol and progesterone at their means plus one standard deviation of the (*ex situ*) population sample. We used the Kolmogorov-Smirnov (KS) test to compare the cumulative distribution functions of estradiol and progesterone across time for the “wait times” events, confirming visual peak pairings [65]. For the KS tests, we conducted (0) lag evaluations on individuals and presented the results at the colony level. We visualized these data to isolate biologically meaningful events versus noise by first comparing data within a given year scale and then subsetting by month. We matched the number of events to plots of peaks per year.

### Assessment of hormones within and among sexes and populations

We tracked progesterone, estradiol, and corticosteroid levels in females. We used linear mixed-effects models to assess individual hormone levels within populations (ACNC, MG, WM), treating individuals as random effects in the R package lme4 [64]. Because progesterone is influenced by estradiol [61], we included estradiol as an additional covariate in the progesterone analysis. We also included season as a random effect. We used log transformations to obtain the “best fit” for the

hormone variables. Log transformation is useful for handling nonparametric data and for visualizing data that spans different scales [65]. We used corrected Akaike Information Criterion in the MuMIn package [66] to select optimal models among fixed or varying slopes and intercepts for covariates.

We used the packages effectsize [67], jtools [68], and sjPlot [69] to derive coefficient estimates, predicted marginal effects, and Nakagawa & Schielzeth’s pseudo-R² of the models [70].

## Results

We obtained n = 865 FHM data points for females housed at the ACNC. We generated FHM data from an additional 77 females from the MG and WM populations, respectively (Table 1). For population-level analyses, the data spanned from April 3, 2015, to August 30, 2019. For *ex situ* ovulation assessment, we analyzed continuous (daily) data from April 3, 2015, to December 31, 2016.

**Table 1.** Red squirrel hormone values obtained from fecal hormone metabolite (FHM) assessment, partitioned by population and sex. The ACNC population is part of an *ex situ* conservation breeding program. Units are ng/g.

| Hormone | Sex | Population | n | Mean | SD | SE | Min | Max |
| --- | --- | --- | --- | --- | --- | --- | --- | --- |
| *Cortisol* | F | Phoenix Zoo | 865 | 53.7 | 60.21 | 0.07 | 1.8 | 1083.8 |
|  | F | Mt. Graham | 31 | 55.4 | 36.10 | 1.16 | 9.8 | 140.9 |
|  | F | White Mountains | 46 | 172.0 | 203.17 | 4.32 | 17.4 | 895.4 |
| *Estradiol* | F | Phoenix Zoo | 810 | 103.4 | 56.37 | 0.06 | 10.9 | 433.4 |
|  | F | Mt. Graham | 31 | 157.9 | 185.03 | 5.97 | 12.1 | 1012.7 |
|  | F | White Mountains | 46 | 376.0 | 595.86 | 12.68 | 25.4 | 3525.0 |
| *Progesterone* | F | Phoenix Zoo | 789 | 27.0 | 10.12 | 0.01 | 3.9 | 73.4 |
|  | F | Mt. Graham | 31 | 84.4 | 87.74 | 2.83 | 12.9 | 317.0 |
|  | F | White Mountains | 46 | 157.2 | 144.69 | 3.08 | 20.9 | 532.3 |

### Hormone compositions among populations

Mean progesterone values were higher in both *in situ* populations compared to the ACNC (Table 1). Mixed models with varying intercepts and slopes for individuals, as well as varying intercepts for seasons, best fit the progesterone data (n = 865, R² = 0.856; Table 2). We found a strong relationship between progesterone and estradiol (p < 0.001), after controlling for other variables and accounting for the random effects of individuals and season. For each 1% increase in estradiol, progesterone increased by 1.9% (95% confidence intervals (CI): 1.7–2.2%; Fig 2). The *ex situ* (ACNC) population differed from both *in situ* populations (*p* <0.001; Fig 3, Table 2); a 1% increase in progesterone at the ACNC population was comparable to a 2.39% (CI: 1.4–4.1%) equivalent increase in MG and 3.13% (CI: 1.9–5.1%) increase in WM populations, respectively.

**Fig 2.**
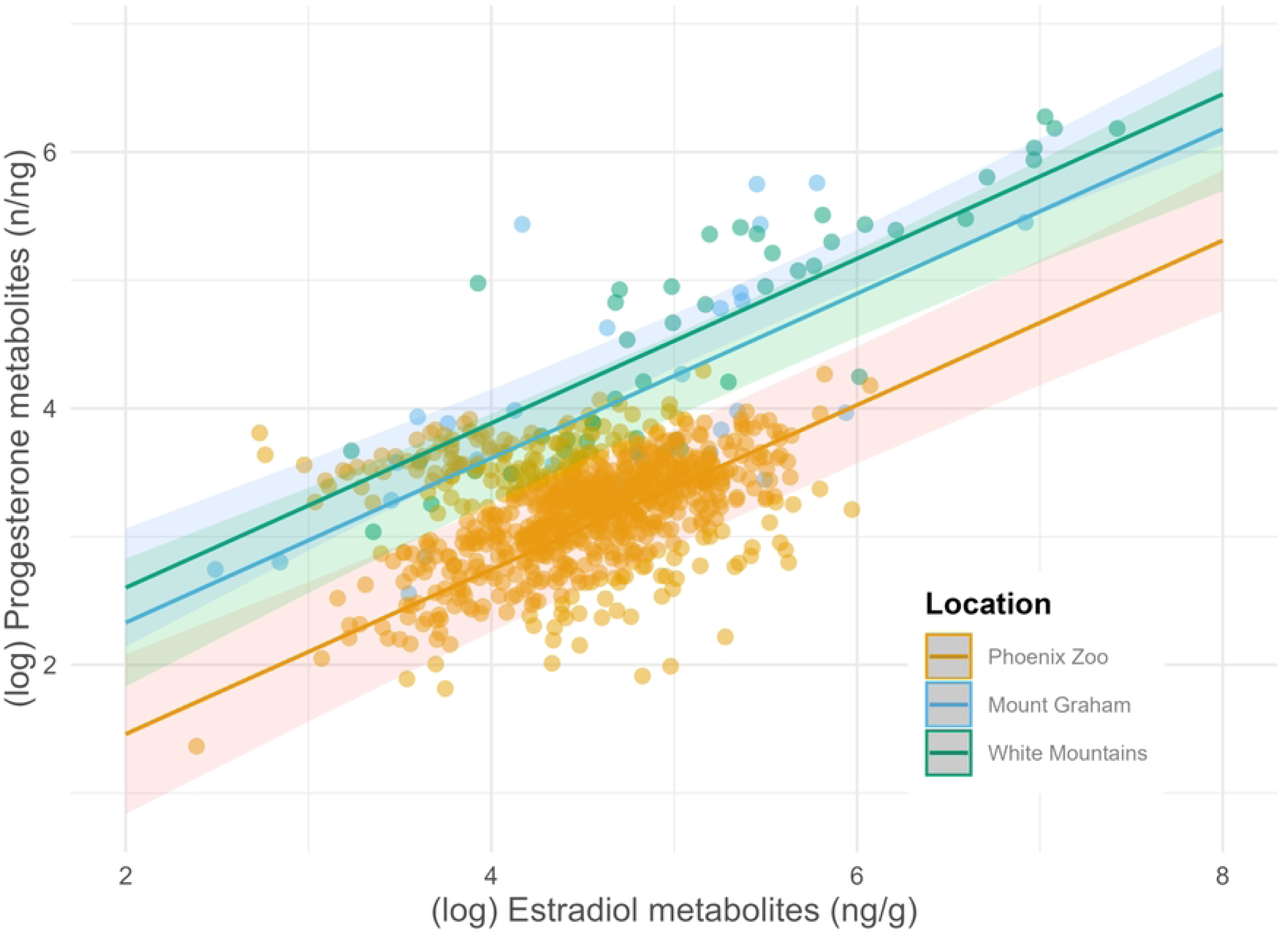
Effects of estradiol on progesterone for three populations of red squirrels, after accounting for random effects of individuals and season. Hormone data are log-transformed, and units are in nanograms per gram. Colors represent population; dark lines are predicted values (marginal effects) for progesterone given other terms; shaded bands are confidence intervals.

**Fig 3.**
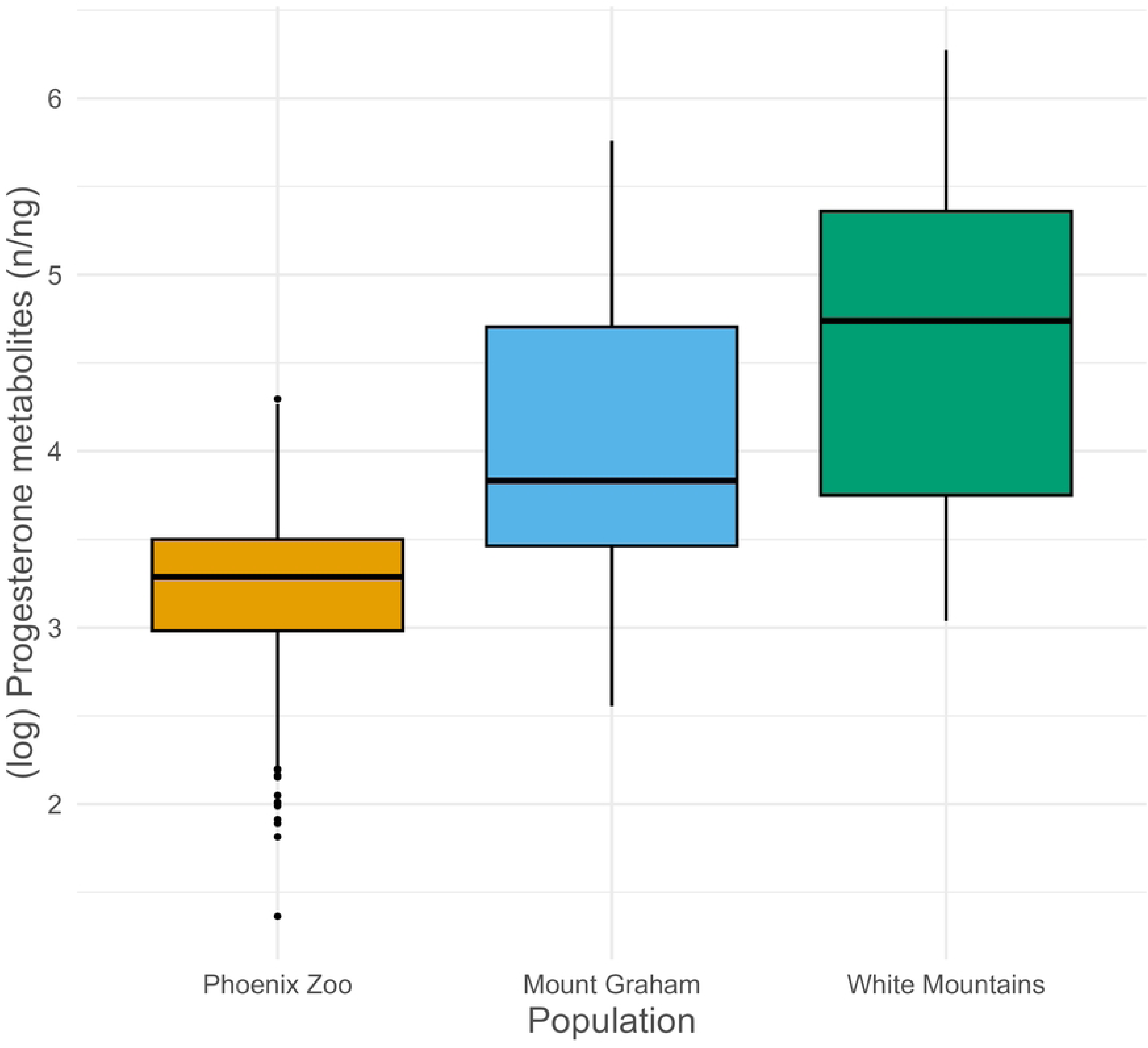
Female progesterone metabolites within three populations of red squirrels in Arizona. Units are nanograms per gram on the log scale.

**Table 2.** Mixed effect models for fecal hormone metabolites (response) in three red squirrel populations in Arizona. Units are ng/g. Bold indicates significance.

| Sex | Response | Predictor | Est | CI | P |
| --- | --- | --- | --- | --- | --- |
| F | Progesterone | Intercept | 1.19 | 0.53–2.69 | 0.674 |
|  |  | Pop: MG | 2.39 | 1.39–4.09 | **0.005** |
|  |  | Pop: WM | 3.13 | 1.94–5.05 | **<0.001** |
|  |  | Est | 1.90 | 1.68–2.15 | **<0.001** |
| F | Estradiol | Intercept | 79.31 | 34.50–182.31 | **<0.001** |
|  |  | Pop: MG | 1.16 | 0.43–3.12 | 0.767 |
|  |  | Pop: WM | 2.43 | 0.99–5.91 | 0.066 |
| F | Corticosteroid | Intercept | 53.56 | 28.99–98.94 | **<0.001** |
|  |  | Pop: MG | 0.92 | 0.45–1.88 | 0.811 |
|  |  | Pop: WM | 2.05 | 1.08–3.90 | **0.046** |

Abbreviations: sex (F = female); Pop = Arizona population (MG = Mount Graham, WM = White Mountains, reference level is *ex situ* population at ACNC).

An estradiol metabolite model was best fit using varying intercepts for individual random effects and omitting season (*n* = 887, *R^2^* = 0.660; Table 2). The *ex situ* and *in situ* MG populations had equivalent estradiol levels, although WM trended towards higher estradiol metabolite levels (*p* = 0.066; Fig 4a.; Table 1).

**Fig 4.**
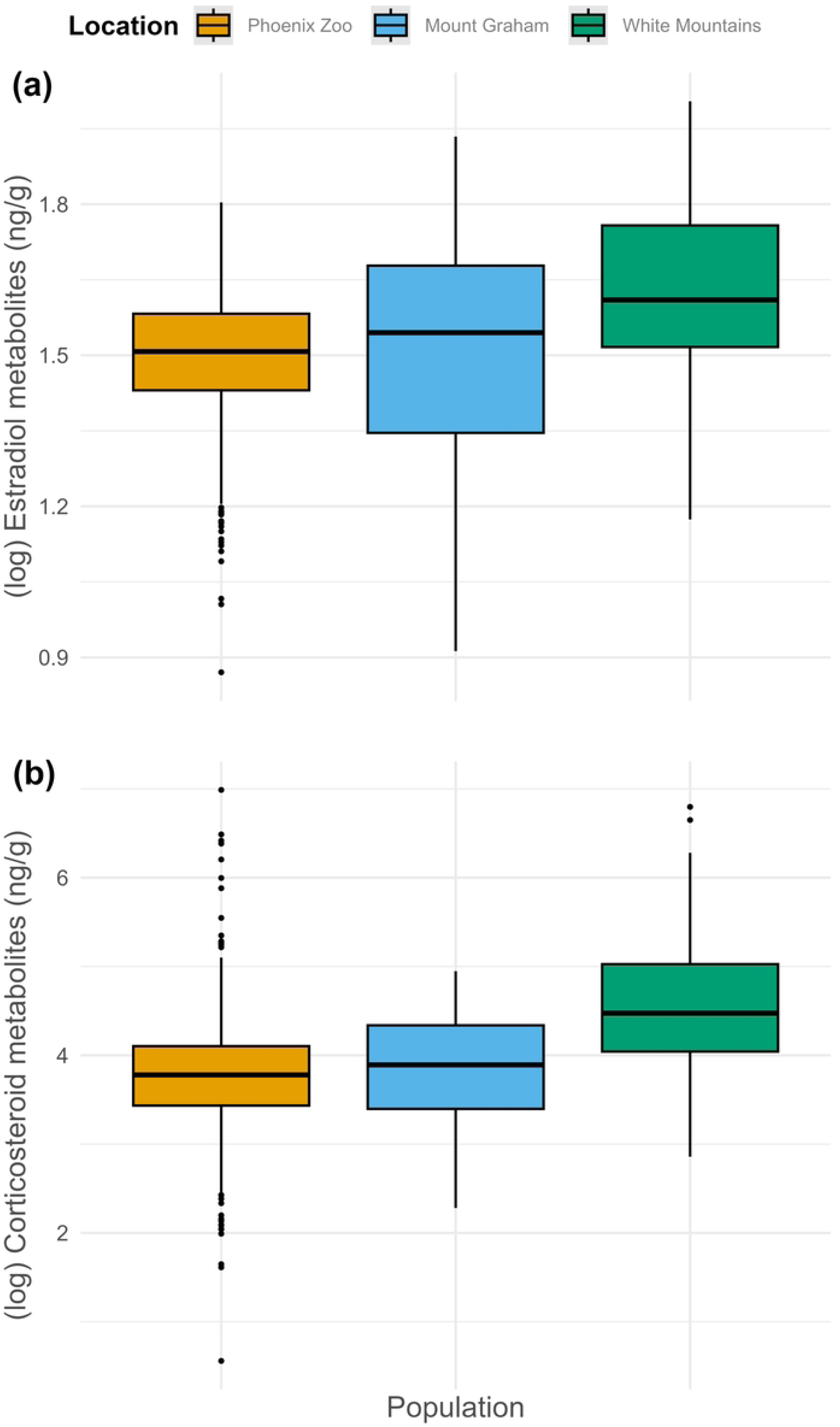
a. Fecal hormone metabolite values for red squirrel populations in Arizona. Hormone metabolites include (a) female estradiol and (b) corticosteroids for females, (Data are colored by population; ACNC is an *ex situ* conservation breeding program. Units are nanograms per gram on the log scale.

A female corticosteroid metabolite model was best fit using varying intercepts for the random effects of individual and season, yielding mixed differences at the population level (n = 955, *R²* = 0.518; Table 2). The WM population yielded a 2.05x (CI: 1.08–3.90x) increase in corticosteroid metabolites compared to the *ex situ* population, after holding other model terms constant (*p* = 0.046). There were no differences, however, between the *ex situ* and *in situ* MG populations for female corticosteroid metabolites (*p* = 0.811; Figure 4b).

### Detecting estrus events

With estradiol–progesterone peak lag set to 0 (i.e., same day), there were six ovulation events in 2015 (mean ± SD time between events: 40.0 ± 58.2 days) (Table 3, Figures 5,6). Notably, events were detected only in May and June. These FHM results support that ovulation in *ex situ* MGRS occurred in regular, spontaneous cycles in 2015 (KS test: *D* = 0.335, *λ* = 0.03, *p* = 0.525). In 2016, we detected 16 events (25.2 ± 28.4 days) (Table 3, Figs 5,6). Notably, events occurred in February, March, May, July, August, October, November, and December. Again, the data support spontaneous ovulation (*D* = 0.246, *λ* = 0.04, *p* = 0.323). These results support the premise that an immediate progesterone peak following a spike in estradiol indicates an ovulation event, and the occurrence intervals over time indicate spontaneous reproductive cycling.

**Fig 5.**
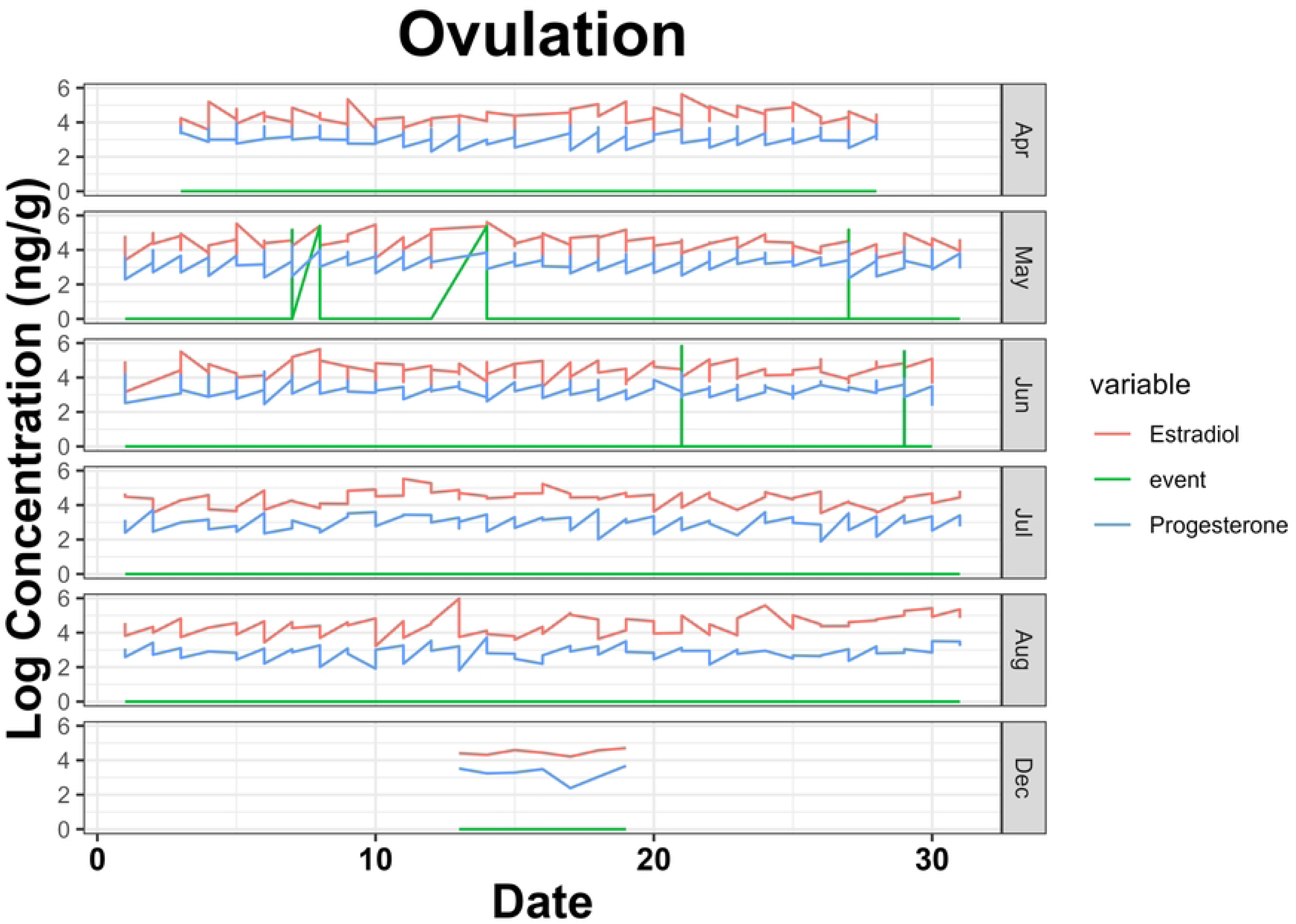
Ovulation events from localmaxima script for Mount Graham red squirrels (*Tamiasciurus fremonti grahmensis)* in 2015. Events (n = 6) were calculated as peaks greater than one standard deviation above the mean for estradiol, followed by progesterone on a 0-day lag. In 2015, events occurred only in May and June.

**Fig 6.**
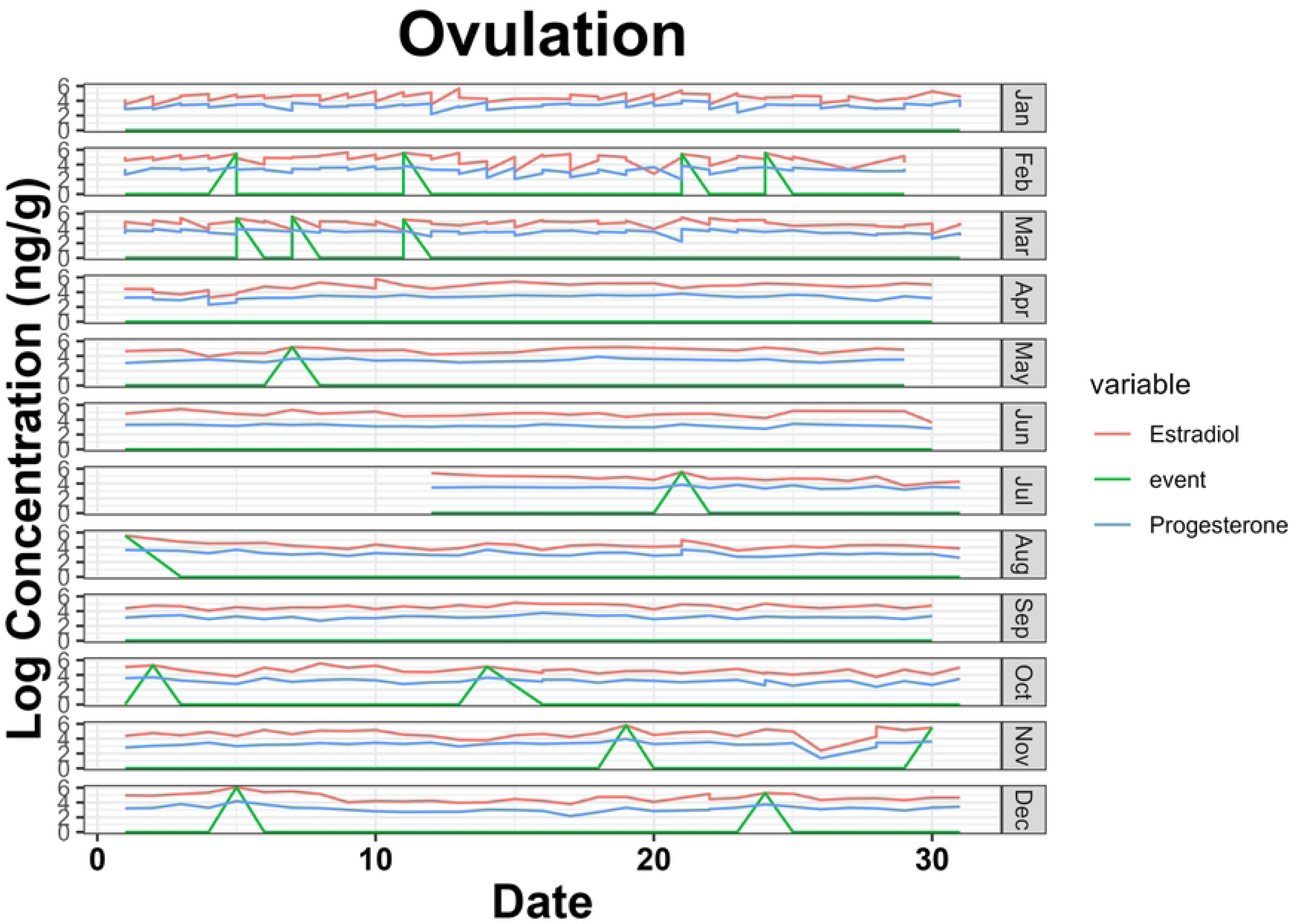
Ovulation events from localmaxima script for *ex situ* Mount Graham red squirrels in 2016. Events (n = 16) were calculated as peaks greater than one standard deviation above the mean for estradiol, followed by progesterone on a 0-day lag. In 2016, events occurred in every month except January, April, June, and September.

**Table 3.** Ovulation events per year in *ex situ* managed Mount Graham red squirrels (*Tamiasciurus fremonti grahmensis)*. Data are results of Kolmogorov-Smirnov (KS) tests of spikes in estradiol followed by progesterone with a 0-day lag (i.e., same day).

| Year | Events (n) | Mean interval (days) | Median interval (days) | SD | *p* |
| --- | --- | --- | --- | --- | --- |
| 2015 | 6 | 40.0 | 19.0 | 58.2 | 0.53 |
| 2016 | 16 | 25.2 | 11.0 | 28.4 | 0.32 |

## Discussion

### *In situ* vs. *ex situ* red squirrel FHM

Our findings revealed that progesterone values among wild populations were similar; however, ex situ MGRS progesterone values were considerably lower than those in *in situ* populations. These findings suggest that *ex situ* progesterone values may be influenced by conditions associated with *ex situ* management or by other tested factors, such as social management [33]. The lower overall progesterone metabolite values of the *ex situ* managed MGRS could be attributed to differences in environmental factors, such as temperature [71], noise [72], and light levels [73,74] as compared to the wild. Alternatively, nutritional factors could be influencing reproductive hormones. Deficiencies in phosphorus, vitamin A, and protein can impact reproduction in livestock [75].

Although we attempted to simulate natural nutrient intake [43] in the *ex situ* MGRS population, we did not evaluate circulating nutrient concentrations within this population. Stress can also impact reproduction [76] and can cause the reduction or cessation of reproductive cycles or spontaneous abortion [33]. Proximity to conspecifics, especially in highly territorial species, can disrupt estrus cycles [33]. Although visual barriers were designed to mitigate visual response cues, auditory and olfactory stimuli may still affect physiological responses [77].

Corticosteroid levels in females were lower in both *ex situ* and *in situ* MGRS than in WM; however, although corticosteroid levels alone may be an insufficient indicator of stress [78], this suggests that at least some aspects of stress are not elevated above normal levels. Additionally, estradiol levels were equivalent between zoo and wild populations. Taken together, it is warranted to investigate the causes of low progesterone while maintaining management practices that ensure other hormones and behaviors are balanced, comparable to wild conditions.

### Estrus detection

MGRS were suspected of having only a single, brief (less than 8 hours) estrus period per year [45,52]. Data indicate that MGRS are likely seasonal breeders, and that ovulation events (polyestry) occur spontaneously during their active breeding seasons. By longitudinally tracking FHM progesterone levels, which increased following elevated estradiol above the mean, we confirmed that MGRS exhibited multiple estrus events over the course of this study, rather than a single estrus event.

We used FHM, an effective noninvasive longitudinal physiological approach to monitor hormonal changes. Noninvasive FHM monitoring provided more accurate, continuous insight into hormone changes over time, underscoring the value of combining physiological data with behavioral inferences rather than relying solely on observational assumptions [79]. This study focused primarily on identifying FHM measures of increases in estradiol accompanied by increases in progesterone as indicators of estrus, applying KS to the occurrence of these events, and characterizing the time between events. Possessing this insight will be a critical complement to behavioral observations, providing an objective, internal perspective on physiological states that may not be apparent from observed behaviors alone and adding a level of quantifiable scientific rigor [32].

We detected 22 ovulation events across two years: 6 in 2015 and 16 in 2016, with a higher rate in the second year, perhaps due to maturity [80]. Our results characterized estrus events by an increase in estradiol followed by an increase in progesterone within a day, and differed appreciably from the Santymire study, which reported detecting three ovulations from a single female over the 18-month study and eight follicular phases collectively [35]. It should be noted that two of the three females in this study were also included in the Santymire study. study, albeit three years later. Upon acquisition in July 2014, these *ex situ* females were estimated to be “young of the year” (mean ± SD: 0.8 ± 0.3 Y) at capture, typically just after dispersing from the nest [49], and it is estimated that maturity occurs approximately seven months after birth [83]. Environmental, physiological, or age factors may also have affected [81]. Further investigation would require tracking metabolite changes in relation to age and external environmental factors, such as light cycles [73,74], noise[72], and nutrient variations [82,83].

Despite lower levels of reproductive hormones, specifically progesterone, analysis of estrus occurrences herein reveals that the *ex situ* population experienced numerous estrus cycles, even when separated from male MGRS, confirming the findings of the Santymire et al. study [35] that MGRS exhibit spontaneous ovulation. However, the number of events detected in this study differed appreciably from their findings. Further study on social, nutritional, environmental, and physical factors of *ex situ* management may provide insights into the reproductive failure in *ex situ* managed MGRS. These revelations make sense; a single annual reproductive event would not be reproductively advantageous for a species with an average lifespan of approximately two years in the wild [45,84,85], primarily due to predation [86]. We believe that a key reason the single-estrus event hypothesis prevailed in the literature is that, absent logitunidal *in situ* FHM monitoring, the reproductive assessment was based mainly on observational data or inductive reasoning, which may be influenced by factors such as limited short-term or intermittent access to animals or highly efficient breeding success, which subsequently result in pregnancy. However, work conducted by Millar [36] concluded, through dissection, that the western red squirrels were likely seasonal breeders, capable of estrus more than once a year.

Applying FHM analyses has facilitated insight into the physiological state of over 100 species since the 1970s [87]. Additionally, FHM analysis has helped identify differences in reproductive\ patterns, the estrous cycle, and reproductive phenology [87], [30]. Noninvasive hormone research has led to a greater understanding of reproductive biology [21], [30] and has revealed insights into the importance of maintaining sociality and/or conspecific spatial distancing in species [33], which can disrupt estrus cycles or prevent them from occurring at all. Our study builds on Santymire et al. [48] by applying FHM for estrus detection, offering advantages over invasive swabbing techniques.

While their work provided insights into MGRS estrus cycles, their approach required collecting vaginal swab samples to detect cornified epithelial cells and measuring vulvar swelling as an estrus indicator. The vaginal swabbing requires some acclimation of the squirrels to swabbing to obtain a sample, and the vulval swelling assessment is determined by visual examination of that area, which may introduce a level of subjectivity. Even minimally invasive methods, such as swabbing, require close human proximity in order to obtain the sample, which can be a stressor and alter natural behaviors, potentially affecting the estrous cycle itself [56], [88]. Each of these studies is an example of the first time estrogen and progesterone FHM analysis was applied to characterize the reproductive biology of a North American squirrel, MGRS. Our approach enables noninvasive collection of FHM data from fecal pellets, providing both noninvasiveness and longitudinally continuous data. However, it is necessary to couple this work with behavioral indices; doing so may mitigate the need for physiological sampling and ultimately reduce or eliminate the need to collect and analyze FHM. This study reveals that reduced fecundity may be due to lower circulating levels of female reproductive hormones, such as estradiol and progesterone [89]. Specifically, progesterone levels were lower in the *ex situ* squirrel population than in both the MG and WM squirrel populations sampled. Progestrone production levels are also highly dependent on light cycles [73] [74,90], and chronically low levels can affect the ability to develop a mature ovum, as it plays a critical role in follicle development [91].

Further research that examines behavioral changes, such as increases or decreases in aggression, alongside changes in estradiol levels, could help refine this work. When combined with behavioral monitoring, FHM can assist reproductive behavior strategies and breeding management development, including methods for determining behaviors indicative of the presence or absence of a physiological reproductive state [92], [27], [93], [94],[17], [95], [96], [97], [98], [99], [100]. Future studies should incorporate a broader range of controlled environmental conditions to identify factors that may influence the observed differences in reproductive physiology between *in situ* and *ex situ* populations, including noise levels, photoperiod, proximity to conspecifics and humans, and temperature.

### Conservation implications and future considerations

Beyond advancing species-specific reproductive knowledge, the results of this study can have direct, actionable implications for endangered species management, particularly for taxa that require *ex situ* assurance populations as part of recovery planning. The demonstrated sensitivity of progesterone metabolites to *ex situ* conditions highlights the need for husbandry protocols that explicitly incorporate physiological monitoring as a routine management tool rather than relying solely on behavioral observations or breeding outcomes. Integrating FHM monitoring into daily or weekly management decisions would allow caretakers to identify suboptimal reproductive states before breeding failure occurs, enabling earlier, more targeted interventions, especially when these data are combined with behavioral indices. Combining FHM estrus characterizations with behavioral indices will ultimately mitigate the need to collect physiological samples, enabling more rapid assessment of reproductive readiness.

These findings underscore the importance of designing *ex situ* environments that more closely reflect species-specific ecological and social requirements. With highly territorial mammals, such as MGRS, spatial arrangement, auditory exposure, and olfactory cues should be treated as core components of reproductive management rather than secondary considerations. The observed progesterone suppression in *ex situ* females suggests that even when estrus and ovulation occur, subtle disruptions in environmental or social parameters may impair luteal function and successful conception [33]. Applying physiological feedback to enclosure design, visual and acoustic buffering, and seasonal adjustments in animal spacing could substantially improve reproductive outcomes for similarly territorial or stress-sensitive species.

The confirmation of spontaneous polyestry through longitudinal, noninvasive FHM monitoring has implications for breeding program timing and genetic management. Rather than concentrating breeding attempts around narrow reproductive windows, managers can use real-time hormone profiles to align pairings with verified physiological readiness. This approach can reduce unnecessary animal transfers, repeated introductions, and handling-related stress, while increasing the likelihood of successful matings and improved genetic representation across breeding seasons. Such precision breeding strategies are particularly valuable for small populations where each reproductive opportunity carries disproportionate importance.

More broadly, this work sets the foundation for coupling longitudinal physiological data with behavioral monitoring to refine species-specific reproductive indicators. Applying similar frameworks across endangered taxa can improve the interpretation of ambiguous behaviors, reduce reliance on inductive assumptions, and support evidence-based decision-making. The scalability of fecal hormone metabolite analysis makes it especially well-suited for diverse conservation applications, including field-based monitoring of reintroduced populations, assessment of translocation readiness, and post-release reproductive evaluation.

Collectively, these applications position noninvasive endocrine monitoring as a cornerstone tool for modern conservation breeding programs[21,101]. By linking reproductive physiology to environmental, nutritional, social, and behavioral variables, this approach enhances adaptive management capacity and supports more resilient *ex situ* populations that align with long-term *in situ* species recovery goals.

## Conclusion

This study meaningfully advances the understanding of MGRS reproductive physiology, particularly by confirming polyestrous cycling through longitudinal hormone tracking, a technique that mitigates the biases introduced by reliance on observational data alone. By employing a noninvasive and stress-mitigating method, our study lays the groundwork for future research on the reproduction of endangered mammals, particularly in *ex situ* conservation settings where little or no reproductive biology is known. As we advance, these findings should inform improved husbandry practices and could be applied to a broader range of species, ultimately supporting global conservation efforts.

## Acknowledgments

We thank the USFWS, Arizona Ecological Services office, the volunteers and staff at the Arizona Center for Nature Conservation/Phoenix Zoo, and the Conservation Research Lab at the University of Arizona. T & E, Inc., the Arizona Game and Fish Department Heritage Fund, and the University of Arizona provided funding and support. We thank Maria Morandani for providing access to WM and MGRS samples and Brian Blais for his assistance with data analysis. We want to thank Rachel Santymire, who headed the Lincoln Park Zoo Davee Center for Epidemiology, and her lab technician, who processed the samples for this study. We wish to thank Samir Rachid Zaim, Ahyoung Amy Kim, Haozhe Xu, and Elmira Torabzadehkhorasani for their assistance in developing the estrus detection model. We want to acknowledge Brandon Meyer and Jeffrey Oliver for their assistance with troubleshooting the estrus detection model.

